# Targeted detection of endogenous LINE-1 proteins and ORF2p interactions

**DOI:** 10.1101/2024.11.20.624490

**Authors:** Mathias I. Nielsen, Justina C. Wolters, Omar G. Rosas Bringas, Hua Jiang, Luciano H. Di Stefano, Mehrnoosh Oghbaie, Samira Hozeifi, Mats J. Nitert, Alienke van Pijkeren, Marieke Smit, Lars ter Morsche, Apostolos Mourtzinos, Vikram Deshpande, Martin S. Taylor, John LaCava

## Abstract

**Background:** Both the expression and activities of LINE-1 (L1) retrotransposons are known to occur in numerous cell-types and are implicated in pathobiological contexts such as aging-related inflammation, autoimmunity, and in cancers. L1s encode two proteins that are translated from bicistronic transcripts. The translation product of *ORF1* (ORF1p) has been robustly detected by immunoassays and shotgun mass spectrometry (MS). Yet, more sensitive detection methods would enhance the use of ORF1p as a clinical biomarker. In contrast, until now, no direct evidence of endogenous L1 *ORF2* translation to protein (ORF2p) has been shown. Instead, assays for ORF2p have been limited to ectopic L1 ORF over-expression contexts and to indirect detection of endogenous ORF2p enzymatic activity, such as by the sequencing of *de novo* genomic insertions. Immunoassays for endogenous ORF2p have been problematic, producing apparent false positives due to cross-reactivities, and shotgun MS has not yielded reliable evidence of ORF2p peptides in biological samples.

**Results:** Here we present targeted mass spectrometry assays, selected and parallel reaction monitoring (SRM and PRM, respectively) to detect and quantify L1 ORF1p and ORF2p at their endogenous abundances. We were able to quantify ORF1p and ORF2p present in our samples down to a range in the low attomoles. Confident in our ability to affinity enrich ORF2p, we describe an interactome associated with endogenous ORF2-containing macromolecular assemblies.

**Conclusion:** This is the first assay to demonstrate sensitive and robust quantitation of endogenous ORF2p. The ability to assay ORF2p directly and quantitatively will improve our understanding of the developmental and diseased cell states where L1 expression and its activity naturally occur. The ability to simultaneously assay endogenous L1 ORF1p and ORF2p is an important step forward for L1 analytical biochemistry. Endogenous ORF2p interactomes can now be presented with confidence that ORF2p is among the enriched proteins.

## Background

Retrotransposons are endogenous DNA sequences that proliferate to new genomic loci through RNA intermediates: the RNAs serve as templates for the synthesis of cDNAs that are inserted back into the host genome (retrotransposition). In humans, Long Interspersed Element 1 (LINE-1; L1) is the only active and autonomous retrotransposon family. L1s are autonomous because their RNA intermediates encode proteins (ORF1p and ORF2p) that possess biochemical properties and enzymatic activities essential for its retrotransposition. Indeed, these same proteins are essential for the production of processed pseudogenes (a.k.a. retrocopies) and the L1-encoded enzyme, ORF2p (an endonuclease [1] and reverse transcriptase [2]), is also essential for the mobilization of active, non-autonomous retrotransposon-types (SINEs, SVAs). Taken together, retro-mobilized sequences in the human genome constitute a large proportion of the DNA (≳ 45%), consisting of nearly two million repeat-derived sequences [3, 4]. The overwhelming majority of retrotransposon-derived genome sequences are degenerate, inactive ‘fossils’ – the consequence of an ancient evolutionary arms race between host genomes and genetic parasites [5–7]. Yet, around one hundred L1 loci containing intact ORF1 and ORF2 DNA sequences are predicted, or have been observed, to be capable of retrotransposition [8–10]. These loci are constitutively repressed by multiple mechanisms in most cell types and circumstances (e.g. reviewed in [11, 12]). Importantly, dysregulated L1 expression and activity is understood to be a hallmark of human cancers [13–15] and may also contribute to autoimmunity [16], aging [17], and other disease states. Recently, we showed that ORF1p has broad potential as a cancer diagnostic using an ultrasensitive ‘digital ELISA’ and selected reaction monitoring (SRM) mass spectrometry (MS) [18]. Yet, the presence of ORF2p in human cancers has only been inferred by the observation that new ORF2p-mobilized DNA sequence insertions do occur (reviewed in [19]; in endogenous contexts, copies of ORF2p have repeatedly proven to be too few to directly observe and quantify at the protein-level by standard means [20–24].

Intact L1s genes encode ORF1p and ORF2p within single, bicistronic RNAs (L1 ORF1p carries the cognate gene symbol L1RE1 in UniProt; L1 ORF2p has not been assigned a recommended gene symbol in UniProt). These proteins have been shown to preferentially bind to the L1 RNA that encodes them (termed cis-preference [25, 26]), forming L1 ribonucleoproteins (L1 RNPs). ORF1p is an RNA-binding protein and nucleic acid chaperone, with homotrimeric and larger oligomeric quaternary structure, that coats the L1 RNA upon self-assembly [27–30]. While ORF1p is translated in the canonical way, the mechanism of ORF2p translation, from the internal position of its open reading frame in L1 RNA, is unusual and apparently inefficient [31, 32]. In contrast to ORF1p, ORF2p is presumed to be present in only one (or very few) copies per RNP. Calculations performed on affinity enriched, ectopically expressed L1 RNPs (where both ORF proteins can be detected and quantified by shotgun MS) estimated the ORF1p:ORF2p stoichiometry at ≲ 30:1 [20, 33]. However, when ectopic and endogenous *α*-ORF1p co-immunoprecipitates (IPs) were compared side-by-side, it appeared that ectopic expression artifactually elevated ORF2p copy number, skewing the apparent ORF1p:ORF2p stoichiometry [24]. Qualitatively, this discrepancy has been known to the field for some time. Hence, the prevailing model posits that endogenous ORF1p is produced in multiple orders of magnitude greater abundance than ORF2p. Lacking a reliable, quantitative readout on endogenous ORF2p, the compositions, stoichiometries, and heterogeneity of L1 RNPs remain open to further elucidation.

Several groups have independently developed antibodies recognizing human ORF2p. Two studies report detection by western blotting, limited to ectopically expressed ORF2p only [21, 24]. These reports align with the expectation that endogenous ORF2p is extremely lowly abundant, and presumably below the Kd of the antibodies in question. Another study reported detection of endogenous ORF2p in malignant tissues [34] - however, off-target cross-reactivity (with BCLAF1) was later demonstrated [23], warranting additional research to fully characterize and understand the reagent. Yet another study reported generating a polyclonal antibody that detects ORF2p [35], but this reagent lacks further essential validation, e.g. by protein MS and/or L1 RNAi coupled with *α*-ORF2p western blotting in an endogenously expressing cell line, leaving its reliability uncertain. A MALDI MS-validated *α*-ORF2p polyclonal antibody has previously been reported, but this reagent has been exhausted and is no longer available [36]. It is especially notable that multiple groups were unable to detect ORF2p peptides in samples subjected to either extensive fractionation, to *α*-ORF1p IP, or to *α*-ORF2p IP, followed by shotgun liquid chromatography (LC)-MS/MS-based profiling [22–24] - this clearly indicates that ORF2p abundance is lower than shotgun MS is able to reliably detect and quantify (e.g. estimated at *∼*1-10 fmol on the LC column depending on the sample complexity [37]). This apparently also holds true for the detection of ORF2p by western blotting using most published antibodies; ORF2p, even with prior affinity enrichment, is below the level of detection. A lower limit of detection of *∼*1-10 pg is often stipulated when using enhanced chemiluminescent detection in conjunction with highly optimized blotting and detection procedures (corresponding to fmol-amol range), although without significant optimization effort ≳ 100 pg detection limit is likely more common [38–40]. We therefore speculated that more sensitive analytical strategies would be required [24].

In the present study we describe two targeted MS assays known as selected and parallel reaction monitoring, respectively (SRM and PRM) [41–43], which are able to detect and quantify ORF1p and ORF2p in various endogenous expression contexts. Targeted MS assays are of great utility for their higher quantitative sensitivity (≳ 10x that offered by shotgun MS [37, 44]), and also their robustness against false positive identifications, which immunoassays lack [41, 45]. In targeted MS, multiple target transitions are monitored, permitting high confidence assignments of the specific chemical identities of chosen reporter peptides. ORF1p could be detected and quantified in cell extracts and IPs while ORF2p could only be detected and quantified after first being concentrated and enriched by *α*-ORF2p IP, consistent with its much lower abundance in cells. Given our ability to validate ORF2p in our IPs, we also report ORF2p-associated protein interactions [46, 47].

## Results

### Endogenous ORF2p eludes reliable, routine detection

We previously compared *α*-ORF1p IPs performed on HEK293T cells ectopically expressing L1 from a plasmid (L1RP expressed from pMT302 [33]) to *α*-ORF1p IPs performed on resected colorectal cancers (CRC) [24]. While both ORF1p and ORF2p were robustly detected within *α*-ORF1p IPs from ectopically expressing cells, only ORF1p was detected in CRC-derived samples. If any ORF2p was present in the CRC-derived sample, it was below the level of detection, despite exhibiting ORF1p levels comparable to ectopic expression (**Fig. 1A**: compare ORF1p and ORF2p signals in the lane “*α*-ORF1p T” to the dilutions series “*α*-ORF1p pMT302”). Since ORF2p was not observed to co-IP with endogenous ORF1p in these trials, we set out to determine if ORF2p could be enriched and detected by direct IP. During the course of initial testing, *α*-ORF2p cl. 9 was deemed to be promising for IP-based experiments, compared to our other available *α*-ORF2p antibodies (e.g., clones, 5 and 11) [24] based on the yield of ORF2p from cell extracts harboring ectopically expressed L1. Additionally, when conducting endogenous *α*-ORF2p IPs using cl. 9, in various embryonal carcinoma cell lines, we observed a readily detectable yield of ORF1p in IP-western blotting trials (e.g., **Fig. 1B**). *α*-ORF2p cl. 9 was therefore moved forward for use in the present study and we conducted multiple endogenous *α*-ORF2p IP-western blotting and IP-MS trials in an effort to enrich and detect ORF2p directly. Whilst signals at *∼*150 kDa (consistent with the mass of ORF2p) were occasionally detected by western blotting, these signals were (1) neither robust nor consistently reproducible, (2) were also present in some negative controls, and (3) were not the most intense signals on the blot (**Fig. 1B**); furthermore, SDS-PAGE-based shotgun (GeLC-)MS of discrete excised bands around 150 kDa mass did not yield ORF2p peptides (data not shown; see also *Endogenous ORF2p interactomics*, below). The presence and enrichment status of ORF2p in endogenous IPs thus remained decidedly ambiguous and reinforced our prior conclusion that more sensitive and robust approaches were needed [24].

**Fig. 1.**
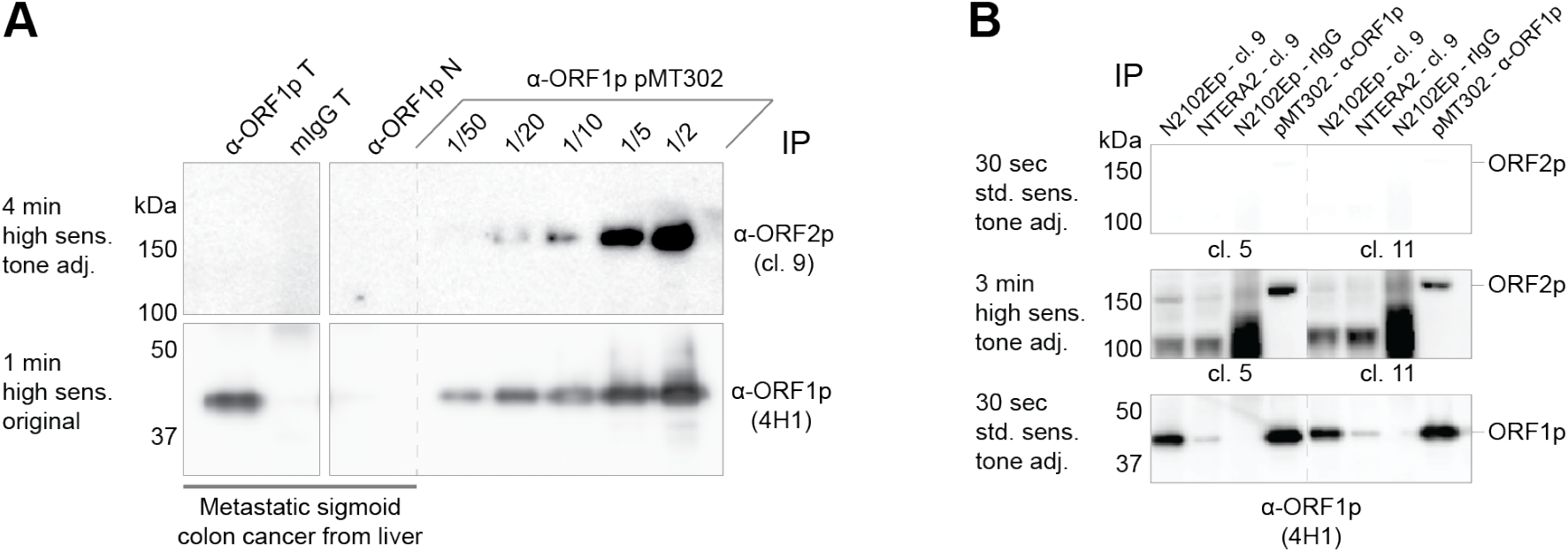
Robust ORF2p signals are absent from IP-western blotting. **(A)** This panel was modified from [24]: *α*-ORF1p IPs were performed on HEK293T cells ectopically expressing L1RP (*α*-ORF1p pMT302), a metastatic sigmoid colon cancer tissue resected from liver (*α*-ORF1p T), and controls (mIgG T: mock IP with naïve mouse polyclonal IgG; *α*-ORF1p N: IP using matched normal tissue). The eluted proteins were probed for the presence of ORF1p (clone 4H1) and ORF2p (clone 9) by western blotting (all samples shown were probed on the same blot). The duration of the image data collection (in min, to the ECL reaction), the CCD imager sensitivity setting (high sensitivity), and whether the panel was tone-adjusted in the figure, are given (see *Methods*). **(B)** This panel displays ORF2p IP-western blot trials to assay the detectability of endogenous ORF2p by direct IP by this method. Enrichment of endogenous ORF2p was attempted using *α*-ORF2p (clone 9) in two embryonal carcinoma cell lines (N2102Ep clone 2/A6 and NTERA-2 clone D1). As a negative control, naïve rabbit IgG was used for IP (rIgG; N2102Ep). As a positive control *α*-ORF1p (4H1) was used for IP (ectopic L1 expression in HEK293T _LD_ from pMT302 [33] which carries ORF2p::3xFLAG). For detection of ORF2p, two different *α*-ORF2p antibodies targeting distinct epitopes were used (clone 5 [left]; clone 11 [right] [24]). For detection of ORF1p, 4H1 was used (as in panel A). ORF2p IPs demonstrated several features that raised doubts about the reliability of the potential ORF2p signal at ∽150 kDa.

### Developing and validating targeted proteomics for the detection of ORF1p and ORF2p

We next turned to SRM (on triple quadrupole mass spectrometer, see *Methods* for assay development details) as a more sensitive and reliable means to detect and quantify both proteins in parallel. For this, we selected a panel of ‘quantotypic’ ORF1p and ORF2p reporter peptides; the selection determinants can be found in **Supplemental Table 1** and the properties of selected peptides meeting the criteria are listed in **Supplemental Table 2**. To validate the assay, we examined its linearity and reproducibility. For linearity, a digest from an extract containing L1 RNPs (expressed from pLD401 [33]) was spiked with isotopically-labeled standards [48] of four peptides targeting ORF1p and ORF2p, respectively, in quantities ranging from 1 amol to 10 fmol (**Supp. Table 3**). Six of the peptide standards show good linearity with the targeted peptides (*R*^2^*>*0.9). The ORF2p peptide TAWYWYQNR exhibited slightly less sensitivity (detection trending toward the fmol range rather than the amol range, *R*^2^=0.83) and the ORF1p peptide LENTLQDIIQENFPNLAR was lacking sufficient sensitivity for detection of the protein in the fmol range; these peptides were therefore excluded for quantitative purposes. To test reproducibility, we analyzed five technical replicates of the extract digestion containing the ectopically expressed L1 at the low end for ORF2p detection (10 amol isotopically-labeled standard) in combination with ORF1p detection. In these measurements, ORF1p was quantified with three peptides (LSFISEGEIK, NLEECITR and QANVQIQEIQR), averaging at 3.1 fmol, and ORF2p was quantified with two peptides (IFATYSSDK and QVLSDLQR), averaging at 10 amol. Coefficients of variation for these peptide quantitations ranged from 10.2-17.2% for ORF1p peptides and 16.9-25.4% for ORF2p peptides. From these measurements, the bulk ORF1p:ORF2p ratio in the cell extract was estimated as 314:1 (**Supp. Table 4**).

To expand the usability of the targeted assay, we also developed a PRM method applicable for the widely-used Orbitrap mass spectrometers. We selected four of the peptides we validated using SRM (ORF1p: NLEECITR, LSFISEGEIK; ORF2p: QVLSDLQR, IFATYSSDK; see **Supp. Table 2**), and tested their linearity using PRM in quantities ranging from 1 amol to 2.5 fmol (**Supp. Table 5**). In agreement with the SRM experiments, we found that these peptides exhibit good linearity (*R*^2^*>*0.9) by PRM within the tested range. To ensure these peptides remained the best options for L1 ORF PRM (allowing for the possibility that the Orbitrap provides different responses for the peptides), we verified the peptide detection across tryptic fragments when using full-length ORF2p as a starting material. For this we used both purified recombinant ORF2p (rORF2p) [49] and *α*-ORF2p IPs from HEK239T cells expressing L1 RNPs from pLD401 [33]. Analyzing a digest of rORF2p in a shotgun MS experiment resulted in a list of 38 peptides that were used to create a targeted PRM method. When both recombinant ORF2p and L1 RNP IPs were analyzed using this method, we found that 24 of the peptides could be detected reliably in both sample types and with a good peak coverage in both MS1 and MS2 (**Fig. 2A & Supp. Table 6**). Based on this broad screen, the QVLSDLQR and IFATYSSDK peptides gave the highest signal, indicating that these peptides are not only quantotypic for ORF2p, but they will also give the highest likelihood of ORF2p detection and quantitation in endogenous samples. A similar approach was used to make a screen of ORF1p peptide responses, using L1 RNP IPs. Here, a total of 8 ORF1p peptides were found (**Fig. 2B & Supp. Table 6**). Along with the two quantotypic peptides NLEECITR and LSFISEGEIK selected for the SRM assay, LTADLSAETLQAR was also detected with a high signal in this screen. However, due to the double arginines at the C-terminus of this peptide, it poses a high risk of missed cleavages (producing instead LTADLSAETLQRR), which would adversely affect the accurate quantification potential of this peptide; it was therefore excluded for the quantification of ORF1p.

**Fig. 2.**
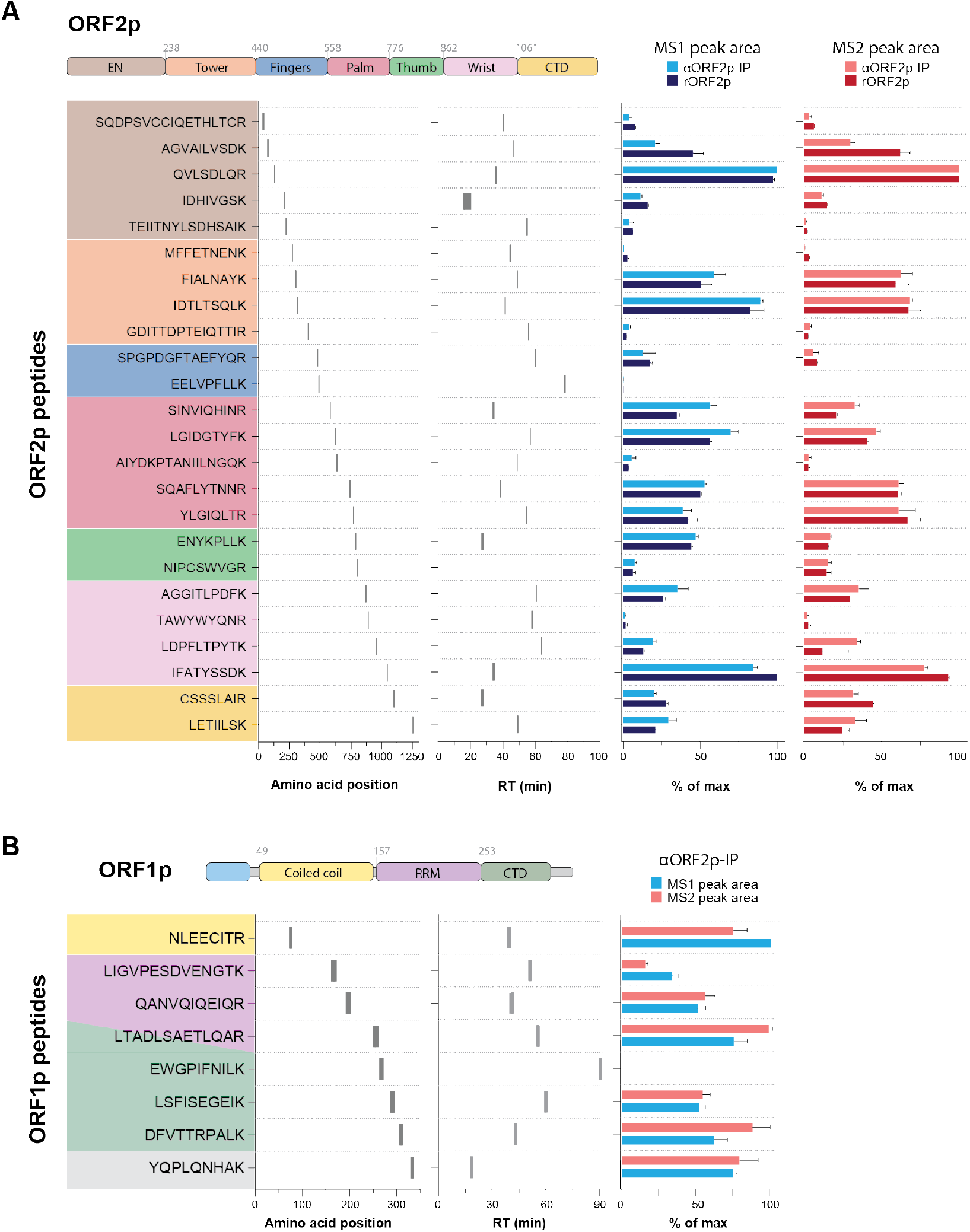
PRM analysis of ORF2p and ORF1p. **(A)** rORF2p (n=2) and *α*-ORF2p-IPs (ectopically expressed L1 RNPs; n=3) were digested with trypsin and all detectable peptides from both samples were analyzed using PRM (targeted precursor m/z values and resulting fragment ions are listed in **Supp. Table 6**). LEFT: all ORF2p peptides are listed from N-terminal (top) to C-terminal (bottom) and colored based on protein domains [49]. Peptide LC retention times are plotted based on peak LC retention times from all sample replicates, using a 90-min gradient from 3-29% solvent B (see *Methods*). Peptide signals are plotted as either summed MS1 precursor peak area from three isotopologues, or summed MS2 peak area from the five most intense fragment ions. Peak areas are represented as % of maximum area observed within each replicate. **(B)** *α*-ORF2p co-IPs of ORF1p (within L1 RNPs, as in panel A; n=3) were digested with trypsin and all detectable peptides were analyzed using PRM (targeted precursor m/z values and used fragment ions are listed in **Supp. Table 6**). The figure layout as in panel A.

### Quantitation of endogenous L1 ORF proteins using SRM and PRM

With both MS assays performing quantitatively down to *∼*10 amol of target analyte injected, we tested their ability to detect and quantify L1 ORFs derived from biological sources that exhibit endogenous expression. For this, we assayed ORF1p and ORF2p peptides (**Fig. 3A**) in cell extracts and in *α*-ORF2p IPs from N2102Ep cells. We successfully detected ORF2p peptides in the IPs but not in cell extracts (**Fig. 3B & Supp. Table 7** for SRM; **Fig. 3C & Supp. Table 8** for PRM): in our current workflow, the IP enrichment step is required for ORF2p peptide signals to rise above the detection limit and/or decrease the signal repression contributed by the high complexity cell extract. We attempted to improve the lower limit of detection for ORF2p in N2102Ep protein extracts by using SDS-PAGE pre-fractionation to reduce the sample complexity and mitigate signal repression (i.e., GeLC-SRM); however, this approach was not successful (data not shown). Averaging the SRM and PRM measurements (*∼*70 amol ORF2p per injection; **Fig. 3D**), and back-calculating (see *Methods*), our data correspond to *∼*20-50 molecules of ORF2p per cell (assuming homogeneous expression of *ORF2* among cells).

**Fig. 3.**
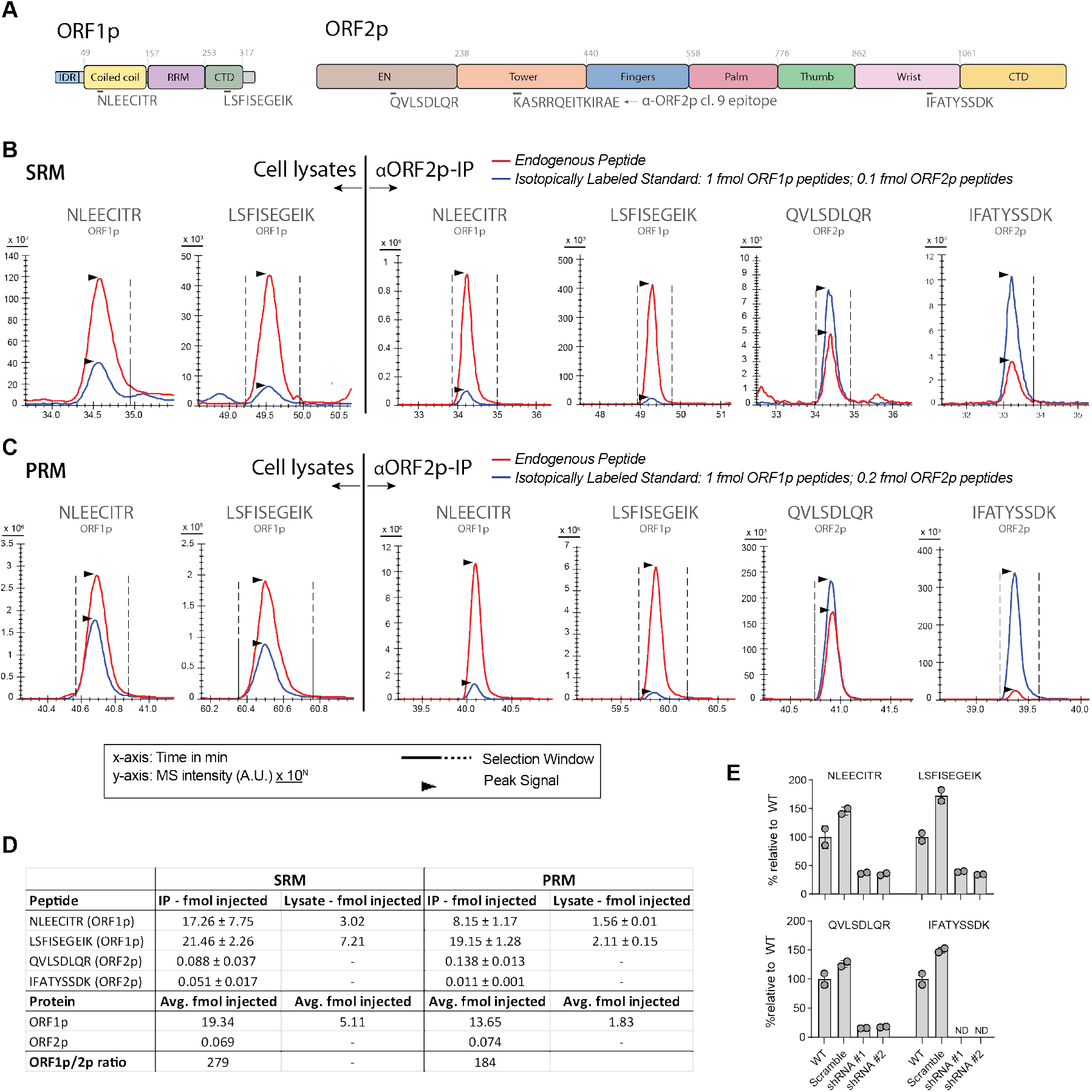
Detection of ORF2p in immunoprecipitates using targeted MS methods. **(A)** Domain structure diagram of ORF1p and ORF2p. The position of the reporter peptides used in the targeted MS assays are highlighted below the diagrams (ORF1p: NLEECITR & LSFISEGEIK; ORF2p: QVLSDLQR & IFATYSSDK). The *α*-ORF2p cl. 9 epitope is also highlighted below the ORF2p structure. **(B)** Detection of L1 peptides in N2102Ep cells using SRM. ORF1p peptides are detected both directly in cell lysates (left) and in *α*-ORF2p-IP (right), while ORF2p peptides are only detected following *α*-ORF2p-IP. Each diagram shows the summed signal of the top-3 fragments from both endogenous peptides (red) and from spike-in of heavy-labeled peptides (blue). The signal from heavy ORF1p peptides corresponds to 1 fmol, and 0.1 fmol for heavy ORF2p peptides. Recorded fragment MS intensities are shown on the y-axis and peptide retention time in minutes are shown on the x-axis. **(C)** Detection of L1 peptides in N2102Ep cells using PRM. Each diagram shows the summed signal of the top-5 (ORF1p) or top-4 (ORF2p) fragments from both endogenous peptides (red) and from spike-in of heavy-labeled peptides (blue). The signal from heavy ORF1p peptides corresponds to 1 fmol, and 0.2 fmol for heavy ORF2p peptides. Recorded fragment MS intensities are shown on the y-axis and peptide retention time in minutes are shown on the x-axis. **(D)** Summary table of results collected for N2102Ep *α*-ORF2p-IPs using either SRM (n=2) or PRM (n=2). **(E)** ORF1p (upper panel) and ORF2p (lower panel) peptide levels detected using PRM-MS in N2102Ep WT cells or cells transfected with either L1-targeting shRNA (shRNA #1 and shRNA #2) or a non-targeting shRNA scramble control. Peptide levels are shown as percent of WT average (n=2). ND: not detected.

Although ORF1p detection for most research purposes can be achieved using standard methods, the targeted MS-based methods allow us to carry out simultaneous quantitation of ORF1p and ORF2p in enriched L1 RNPs and cell extracts (**Figs. 3A-3C**). Using the quantities of ORF1p and ORF2p observed in our *α*-ORF2p IPs, we estimated a molar ratio in the range of *∼*279:1 (SRM) to *∼*184:1 (PRM) in the affinity enriched fraction (**Fig. 3D**); this is approximately an order of magnitude difference from our prior estimates of L1 ORF stoichiometry from ectopic systems (≲ 30:1 [33]). However, we do not assert that the endogenous measurements are indicative of a homogenous L1 RNP stoichiometry (see *Discussion*).

While targeted MS provides high confidence in the chemical identity of the peptides being sequenced, it does not guarantee that those peptides have derived from the presumed target protein (i.e., as opposed to a distinct protein containing an identical peptide). To control for the possibility of false positive ORF2p detection, and further verify the specificity of our assay, we modulated ORF2p abundance in N2102Ep cells by RNA interference knockdown (KD) of L1 RNAs and explored our ability to quantitate the change. For this, using previously validated shRNA sequences [9], we created two independent N2102Ep-derived cell lines (see *Methods*) that down-regulate L1 ORF protein abundances by KD, along with a cognate non-targeting ‘scramble’ shRNA control cell line. We observed that KD by either targeting shRNA reduced the yield of ORF2p peptides observed by IP-PRM compared to WT (unmodified cells) and scramble shRNA control cells (**Fig. 3E**). Averaging the results of the two KDs, we observed a 64% reduction in ORF1p (based on two peptides) and an 83% reduction in ORF2p (based on one peptide), compared to WT; compared to scramble shRNA control cells we observed a 77% reduction in ORF1p (based on two peptides) and an 87% reduction in ORF2p (based on one peptide). The IFATYSSDK peptide of ORF2p was not detected upon KD.

### Quantitation of endogenous L1 ORF2p protein from resected patient tumors

Immunohistochemical staining for ORF1p revealed that it is up-regulated in approximately half of all human cancers [13, 18] and ORF1p has been posited as a promising circulating biomarker for e.g., ovarian and colorectal cancers [18]. Given the elevated levels of ORF1p observed in these tumor-types, and the observation that (presumably ORF2p-driven) L1 insertions accumulate broadly in cancerous tissues [15, 50–53], we hypothesized elevated levels of ORF2p that correlate with those of ORF1p. To test this hypothesis, we applied IP-PRM to resected tumors from patients with either ovarian cancer or colorectal cancer. As expected, all the tumor tissues analyzed were ORF1p-positive, as determined by *α*-ORF1p IP-western blotting (**Fig. 4A**). In the cases where adjacent healthy tissue was resected together with the cancer, these tissues were included in the analyses and also, as expected, were ORF1p-negative.

**Fig. 4.**
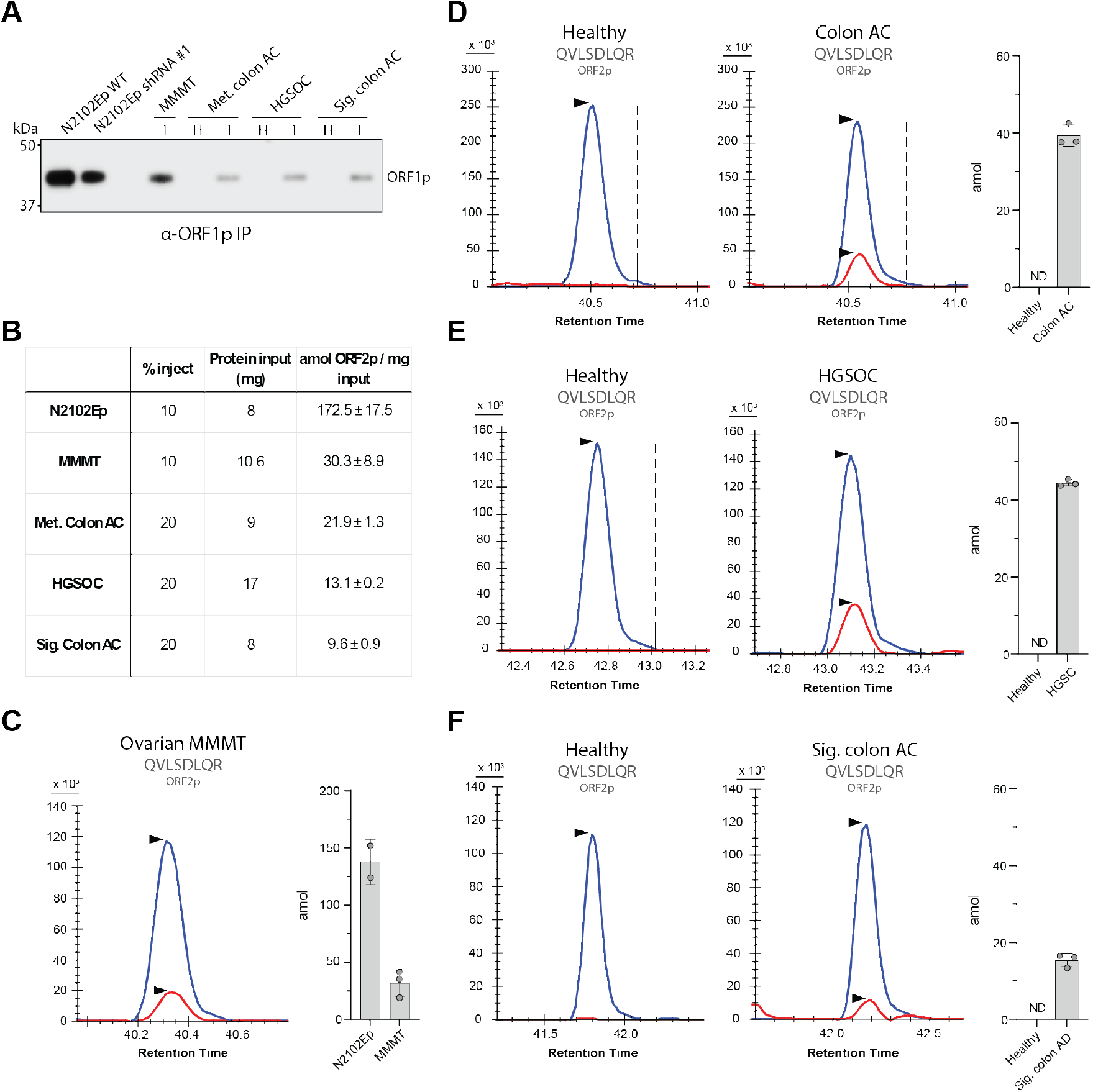
Detection of ORF2p in immunoprecipitates from human cancers. **(A)** *α*-ORF1p western blotting of *α*-ORF1p IPs from N2102Ep WT cells, L1 KD cells, and selected tumors (T) with matched healthy controls (H). **(B)** Table summarizing the quantities of ORF2p (IP-PRM of peptide QVLSDLQR) present in the samples that were blotted in panel A. The ‘%Inject’ column lists the percentage of the sample that was injected into the MS instrument. The ‘Protein Input’ column shows the quantity of total protein used as input for the *α*-ORF2p IPs. The last column shows the detected amount (in amol) of ORF2p adjusted for % injected and the amount of total protein used as the input for the IP. **(C)** PRM quantitation of ORF2p (peptide QVLSDLQR) in *α*-ORF2p-IPs from an ovarian malignant mixed Mullerian tumor (MMMT). The detected levels (in amol) of the peptide, QVLSDLQR, were measured using 10% of *α*-ORF2p-IPs from both N2102Ep cells (n=2) and from the ovarian MMMT patient sample (n=3). In panels C-F, the top-4 fragments from both the endogenous peptide (red) and spiked-in, heavy-labeled peptide (blue) are shown; the heavy peptide corresponds to 200 amol. **(D)** PRM quantitation of the ORF2p in *α*-ORF2p-IPs from a metastatic colon adenocarcinoma (colon AC) resected from the liver. Healthy control tissue corresponds to non-cancerous liver tissue resected together with the cancer. The detected levels of the peptide were measured using 20% of *α*-ORF2p-IPs from either healthy liver tissue (n=3) or metastatic colon adenocarcinoma (n=3). In panels D-F, ND: not detected. **(E)** PRM quantitation of the ORF2p in *α*-ORF2p-IPs from a high-grade serous ovarian carcinoma (HGSOC) tumor. Healthy control tissue corresponds to non-cancerous spleen tissue resected together with the cancer. The detected levels of the peptide were measured using 20% of *α*-ORF2p-IPs from either healthy liver tissue (n=3) or metastatic colon adenocarcinoma (n=3). **(F)** PRM quantitation of the ORF2p in *α*-ORF2p-IPs from a sigmoid colon adenocarcinoma. Healthy control tissue corresponds to adjacent non-cancerous colon tissue resected together with the cancer. The detected levels of ORF2p were measured using 20% of *α*-ORF2p-IPs from either healthy liver tissue (n=3) or metastatic colon adenocarcinoma (n=3).

Using our targeted MS assay, we were able to quantify ORF2p levels from the ORF1p-positive tumors (results summarized in **Fig. 4B & Supp. Table 9**), which included a malignant mixed Mullerian ovarian tumor (MMMT) (**Fig. 4C**), colon adenocarcinoma metastases removed from the liver (**Fig. 4D**), a high-grade serous ovarian carcinoma (HGSOC) (**Fig. 4E**), and a sigmoid colon adenocarcinoma (**Fig. 4F**). In all cases, only the QVLSDLQR peptide could be used for quantitation as the IFATYSSDK peptide levels were below the quantitation threshold. A consequence of this is that quantitations in the tumor samples are based only on the more sensitive QVLSDLQR peptide and the measured moles of ORF2p will therefore appear higher than if using both peptides. Similarly to what was observed for ORF1p, none of the co-resected control tissues had any detectable level of ORF2p. In the MMMT sample, which appeared to have the highest ORF1p and ORF2p levels, ORF2p were measured at 32.1 *±* 9.4 amol which is about four times lower than the amount observed in N2102Ep cells (**Fig. 4C**).

### Endogenous ORF2p interactomics

Having unambiguously demonstrated the presence of ORF2p in our IPs by targeted MS, we sought to describe and report the first endogenous ORF2p-associated interactome. For this, we conducted label-free quantitative shotgun IP-MS analysis of *α*-ORF2p cl.9 IPs, compared with mock IP controls, in N2102Ep cells (**Fig. 5A & Supp. Table 10**). This experiment revealed 487 interactors that were enriched in the IP. However, the mock IP does not control for spurious interactions that form with the target protein or the targeting antibody used for capture [24, 54–56]. This is an important consideration in general [57], but notably important here because ORF2p is likely present in N2102Ep cell extracts at a concentration well below its K_d_ [24]. This can induce poor apparent selectivity of the affinity medium and contribute to the enrichment of more abundant secondary targets by *α*-ORF2p cl.9 paratopes. Our results revealed that the protein L1TD1 was consistently among the most abundant in these *α*-ORF2p IPs. Being that this protein was exapted from L1 within mammalian genomes [58], we explored the possibility that it may contain an epitope with affinity toward the *α*-ORF2p cl.9 antibody. A BLAST search of the presumed antibody target epitope in L1 ORF2p ([333] KASRRQEITKIRAE [346]) [24] revealed this to be the case, identifying a close partial match in the L1TD1 epitope ([501] KASRRQKEI [509]). This raised the possibility that our *α*-ORF2p IP-based interactome was affected by cross-reactivity with L1TD1 (and its associated interactors; see *Discussion*).

**Fig. 5.**
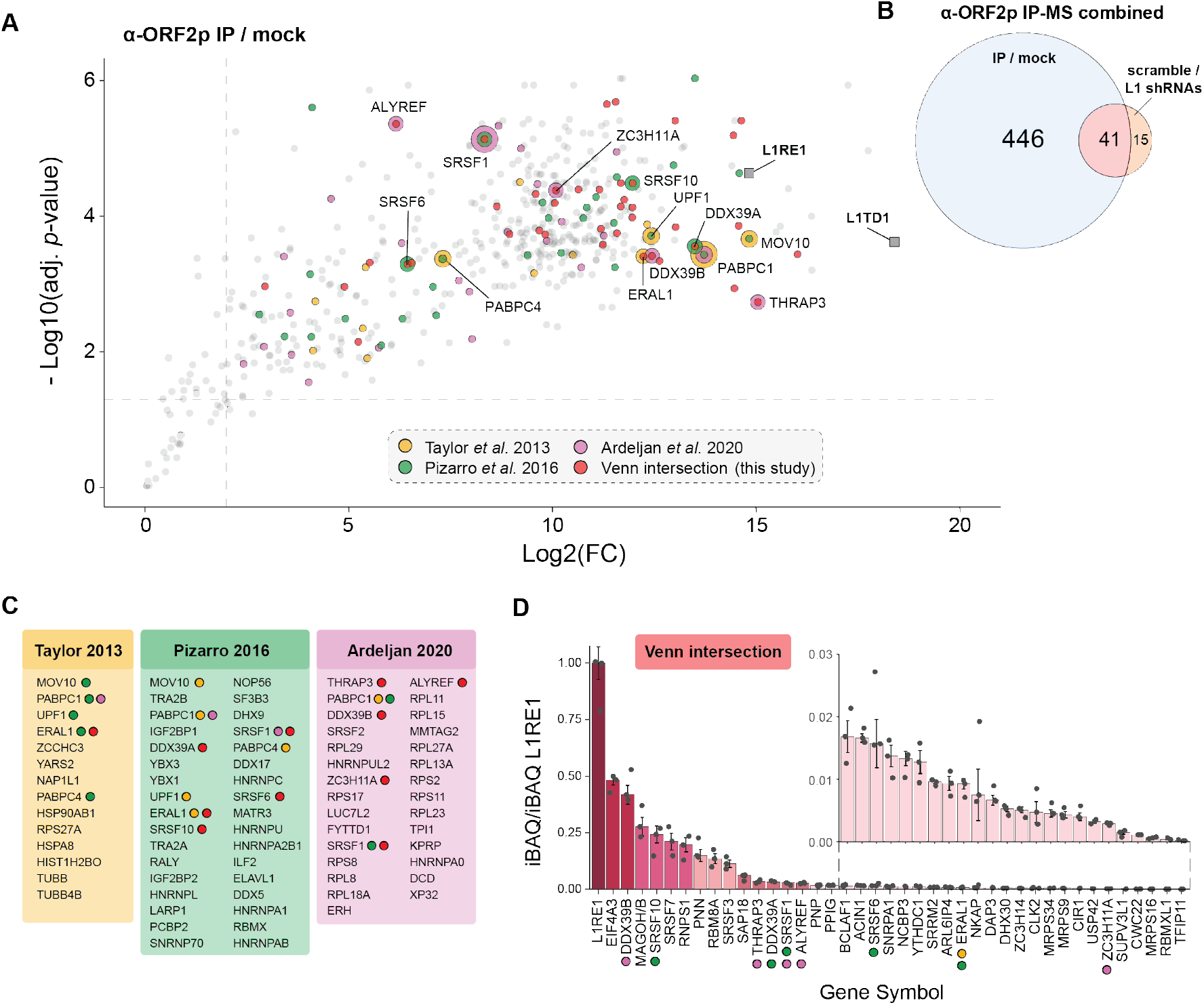
L1 ORF2p interactome in N2102Ep cells. **(A)** The plot displays a one-tailed t-test comparing proteins obtained from label-free quantitative shotgun IP-MS (*α*-ORF2p IP / mock IP; x-axis: log2 fold change; y-axis: -log10 adjusted p-value). Subsets of significant proteins are colored according to their overlap with at least one selected prior published list (described in the legend [12, 24, 33]) and/or their correspondence with a second label-free quantitative shotgun IP-MS analysis from this study (*α*-ORF2p IP scramble shRNA / *α*-ORF2p IP L1-targeting shRNAs; overlap displayed in panels B and D). Proteins that are labeled with text (cognate UniProt gene symbol) correspond with at least two of these lists and display multiple colors accordingly. L1RE1 and L1TD1 are both marked as gray squares. **(B)** Venn diagram showing the number of significant proteins obtained by label-free quantitative shotgun IP-MS: *α*-ORF2p IP / mock IP (DDA) & *α*-ORF2p scramble shRNA / *α*-ORF2p L1-targeting shRNAs (DIA; see *Methods*). **(C)** Lists of proteins that were considered significant in prior publications (as labeled) that also overlap with this study. When a protein was considered significant in multiple prior studies and in this study, colored dots indicate the overlap according to the legend shown in panel A. Note: the significant proteins listed from ‘Pizarro 2016’ were collated and summarized data from two prior publications [59, 60]). **(D)** The plot displays the estimated copy number, relative to L1RE1 (L1 ORF1p), of each of the intersecting 41 proteins in common between the two different IP-MS analyses in this study (panel B). Relative copy number was calculated using the iBAQ values [61] for the proteins listed.

To address this issue and enforce greater signal-to-noise discrimination in our IPs, better controlling for off-target paratope binding, we carried out another series of *α*-ORF2p IPs; this time, we used the same L1-targeting shRNA cell lines and scramble shRNA control cell line, as used in our PRM assay. This experiment revealed 56 proteins that were significantly enriched in *α*-ORF2p IPs conducted using the ‘scramble’ shRNA control cell line (i.e. are associated with ORF2p; **Supp. Table 11**). Importantly, while ORF1p yield was substantially higher in the scramble IP (6.9 fold change), L1TD1 was found to be comparably abundant in both scramble and L1 shRNA IPs – indicating that its yield is not affected by changes in L1 ORF proteins (**Supp. Table 11**). Hence, this experiment allowed us to partially uncouple the L1TD1 associated protein interactions from the ORF2p associated protein interactions in the IPs. Of the 56 proteins that were significantly enriched with ORF2p in scramble IP / L1 shRNA IP comparison, 41 of those were also statistically significant in our IP / mock IP comparison (**Fig. 5B**). A summary of the combined results, including comparisons with other published co-IP experiments, is presented throughout **Figure 5A-D**. When considering only the 41 proteins found to be statistically significant in both the traditional *α*-ORF2p IP and L1 KD *α*-ORF2p IP experiments, we found that the most abundant of these, relative to ORF1p, were proteins associated with exon junction complexes (**Fig. 5D**; see *Discussion*).

## Discussion

### Endogenous L1 ORF protein capture and quantitation – successes and caveats

ORF1p is often expressed in abundance under diseased circumstances in target cells, whereas endogenous ORF2p rarely (if ever) rises to the level of detection of standard proteomic methods (i.e., immuno-assays and shotgun MS), even with prior enrichment from cell extracts (textbfFig. 1). While enrichment may not always be necessary for ORF1p proteomic analysis, the assays we report here provide the opportunity to use affinity enrichment by IP followed by targeted MS (**Fig. 2**) for cases where the most sensitive detection is required (**Figs. 3 & 4**). Our observation is that robust proteomic analysis of endogenous ORF2p requires prior enrichments from cell extracts. Notably, we estimate ≳ 3 orders of magnitude between endogenous ORF1p and ORF2p in N2012Ep cell extracts (see **Fig. 3D**, compare ‘lysate’ levels of ORF1p [fmol] to ORF2p, which is below our limit of detection [amol]). In the literature, antibody-based (immuno) enrichment prior to targeted mass spectrometry is often referred to as iSRM/MRM/PRM [62–65]. These kinds of assays are of clinical importance due to their ability to circumvent limitations of traditional immuno-assays (e.g., ELISAs, blots, etc. [66, 67]). MS offers the unique ability to verify whether the assay is reading-out on the expected analyte, based on several attributes of the data, including chromatographic behavior, fragmentation pattern and accurate mass. Recently, we have used an *α*-ORF1p affinity medium in conjunction with our SRM assay, described in detail here, to validate the performance of a digital ELISA-based ORF1p blood biomarker assay [18]. In the present research we use new ORF1p and ORF2p targeted assays (now developed as SRM and PRM), in conjunction with *α*-ORF2p cl. 9 coupled affinity medium for enrichment [24]. Several *α*-ORF2p antibodies reported in the peer-reviewed literature have been shown to exhibit poor specificity and/or poor selectivity (off-target binding) in cell extracts, in-part because of its low abundance [21, 23, 24]. Indeed, these kinds of effects are common for many antibodies, especially at low antigen concentrations in complex proteinaceous samples [68, 69]. These features also apply to the *α*-ORF2p cl. 9 antibody we have used here [24], e.g., with respect to L1TD1 binding (textbfFig 5 & textbfSupp. Table 11); yet, our affinity medium exhibits sufficient selectivity to enrich ORF2p in the presence of a highly abundant competing epitope. Notably, we validated endogenous ORF2p as a bona fide target of *α*-ORF2p cl. 9 by shRNA KD of L1 in N2102Ep cells (textbfFig. 3E); ORF2p peptide signals are also absent from normal tissues but present in various cancer tissues (**Fig. 4C-F**). This represents, to our knowledge, the first unambiguous, validated demonstration of endogenous ORF2p present in model cells and cancer tissues. It is therefore our contention that no claims of antibody-based endogenous ORF2p detection that lack robust secondary validation, e.g. by MS-based peptide sequencing, should be believed; and here we report requisite targeted assays, with sensitivity down to *∼*10 attomoles, on two common instrument architectures (triple quadrupole for SRM and quadrupole-Orbitrap for PRM), providing in-roads for most MS labs and resource centers.

Although endogenous ORF1p has been less challenging to detect, we have developed equivalent assays for it, in order to simultaneously quantify both proteins. During the preparation of this research, another targeted proteomics assay for ORF1p was published [70]. We note that our selected peptides only included one in common with that study (LSFISEGEIK), the other peptides did not pass our quantotypic criteria due to the risks on missed cleavages (LTADLSAETLQAR and NEQSLQEIWDYVK) and the risks of modifications due to NG-deamidation (LIGVPESD-VENGTK) that could affect accurate quantification. For ORF1p our best results were obtained with the two peptides, NLEECITR and LSFISEGEIK, and for ORF2p our best results were obtained with the two peptides, QVLSDLQR and IFATYSSDK. We note, the N-terminal glutamine in QVLSDLQR may spontaneously convert to pyroglutamate. In our study, we detected this conversion but the ratio of non-converted to converted was comparable for both the endogenous peptide and the heavy-labeled spike-in peptide standard; quantitation using this peptide was therefore not compromised. We observed a difference in quantitation of only *∼*2% when using only the non-converted signal vs. summing both signals.

In principle, our method can detect and quantify both endogenous L1 proteins in parallel, yielding their stoichiometry within L1 RNPs, using affinity-enriched fractions. In the present study we estimated an endogenous ORF1p:ORF2p molar ratio in the *α*-ORF2p enriched L1 RNPs to be in the range of *∼*279:1 (SRM) to *∼*184:1 (PRM). However, we cannot assert this as a bona fide stoichiometry, principally because of the co-enrichment of the protein L1TD1 on the *α*-ORF2p affinity medium we used (Fig. 5 and discussed below in Shotgun Interactomics). L1TD1 is evolutionarily derived from L1 and has been shown to bind L1 RNA and is considered a putative ORF1p interactor [71]. The presence of this protein, if enriched independently from ORF2p in the same IP due to insufficient selectivity, would likely bias the ORF1p quantitation with respect to the fraction exclusively bound within bona fide ORF2p-containing L1 RNPs.

L1TD1 is believed to be directly domesticated from L1 ORF1, however, to our knowledge, no direct relationship has ever been shown between L1TD1 and ORF2p [58]. Notably, neither BLAST nor Smith-Waterman alignments of intact ORF2p against L1TD1 produced any obvious tracts of significant similarity between the two proteins, even failing to align the described ORF2p epitope with the candidate L1TD1 epitope. We only identified the putative L1TD1 epitope alignment by BLAST searching the ORF2p epitope in isolation from other ORF2p as a whole; we believe this leave open the possibility of other hidden, ORF2p-like properties of L1TD1, subject to further investigation. It should be noted that while L1TD1 is expressed in undifferentiated stem cells [72, 73] (including embryonal carcinomas [74] such as N2102Ep cells), it is not apparently ubiquitous in cancer tissues. Yet, it has been reported as a prognostic marker for colon cancer and may be important to cancers that rely on cancer stem cell-like characteristics [75–77]. Hence, reliable stoichiometric estimates of L1 ORFs within enriched RNPs could be made using the *α*-ORF2p cl. 9 antibody and presented targeted methods, provided that no substantial confounding off-target antibody binding to L1TD1 (or other L1 RNP constituents) is observed.

### Quantitation of global ectopic L1 ORF expression

Because ORF2p is highly over-produced in ectopic contexts, ORF2p is detectable by more standard means in these models [20, 24] and targeted MS can be employed to directly measure both L1 ORF proteins in these cell extracts. We therefore used our method to report an estimate of the hitherto undefined stoichiometry of L1 ORF ectopic expression, assayed from cell extracts (from HEK293T_LD_/pLD401 [33]); our measured ORF1p:ORF2p ratio was estimated as 314:1 (**Supp. Table 4**). Users of this assay will likely find that the estimate of ectopic expression stoichiometry varies based on the expression vector genetic context, copy number, cell type, and efficiency of protein extraction (among other possible variables and confounders). Yet, targeted MS measurements can be used to calibrate the performance and set their expectations from ectopic L1 expression laboratory models. Notably, despite the ‘L1 negative’ status of e.g. HEK and HeLa cells that are commonly used for ectopic expression, these cells do exhibit low but detectable levels of endogenous L1 ORF1p [9, 24] – representing a potential quantitative confounder even in these models (although likely to represent only a small fraction of total ORF1p).

### Shotgun Interactomics

Targeted MS confirmed the presence of ORF2p in our IPs (**Figs. 3 & 4**). Yet, L1TD1 is among the most enriched proteins (**Fig. 5 & Supp. Table 10**) and our subsequent interrogation its sequence revealed a candidate target epitope for *α*-ORF2p cl. 9. Being vastly more abundant in the mixture, L1TD1 is likely contributing the majority of co-IP signals. Indeed, we verified that L1TD1 binds to our affinity medium independently from ORF2p by shRNA KD of L1 (which diminishes the levels of both ORF1p and ORF2p; **Fig. 3E**): L1TD1 accumulation on the affinity medium was unaffected by this manipulation (**Supp. Table 10**), supporting the notion that most of this protein is bound to *α*-ORF2p cl. 9 affinity medium independently from L1 ORF proteins. Yet, the matter is complicated by the fact that L1TD1 has been characterized as a putative L1 RNP constituent [71]. To avoid polluting our interpretation of ORF2p-interacting proteins with proteins contributed independently through interactions with L1TD1, we filtered our statistically significant hits to meet at least one of two additional criteria: (1) they were previously characterized in a published L1 interactome study in e.g., HEK cells, HeLa cells, or colorectal cancers (where different antibodies were used for enrichment); (2) they were statistically significant in both the *α*-ORF2p IP / mock IP and *α*-ORF2p scramble shRNA IP / *α*-ORF2p L1-targeting shRNA IP comparisons (**Fig. 5**). Post-filtering, several previously defined high-confidence and functionally validated L1 RNP interacting proteins were retained, including e.g., MOV10, PABPC1, PABPC4, UPF1, and ZCCHC3 [33, 78–80]. The candidate ORF2p interactome we presented here also revealed that exon junction complexes (EJCs, i.e., MAGOH, RBM8A, EIF4A3) and multiple proteins functionally-linked to pre-mRNA splicing, EJCs, and/or mRNA export are present (e.g., highlighted in **Fig. 5D**), co-enrich with ORF2p, independently of L1TD1. Indeed, we estimate EIF4A3 and DDX39B are approximately one-half as abundant as ORF1p in the *α*-ORF2p IPs. We speculate that the co-enrichment of these EJCs reflects an abundance of mRNAs present in the same heterogenous macromolecular assemblies as L1 RNPs [46, 47, 81] in N2102Ep cells. Based on our previous studies in HEK293T, we have speculated these assemblies resemble IGF2BP1 (IMP1) granules - which are known to enrich mRNAs, EJCs, and related proteins, among others [78, 82].

We are cautious not to over-interpret these findings and instead consider them a proof of concept that endogenous ORF2p interactomics are now possible, with rigorous validation provided by the targeted MS assays presented. We successfully used RNA interference KD of L1 RNA to uncouple L1 ORF protein interactors from L1TD1 interactors, yet the statistical significance of our hits and their according effect sizes will be mitigated by the penetrance of the KD (*∼*80-90% compared to scramble, considering both ORF1p and ORF2p). Untangling the contributions of L1TD1 to L1 RNPs requires further research.

## Conclusion

The appearance of endogenous L1 ORF proteins within cellular proteomes is typically associated with dysfunction and disease, motivating methods to (1) quantitate their presence (e.g., as biomarkers) and (2) understand their potential pathological contributions e.g., via induced alterations in macromolecular networks and responses linked to DNA damage and inflammation. While both proteins are generally absent or of low abundance in most cell-types during healthy homeostasis, ORF1p reaches analytically tractable levels in many disease states; yet, due to its orders of magnitude lower abundance, ORF2p has eluded robust and reliable detection and quantitation in model cells and patient samples. Our targeted MS-based assays solve L1 ORF detection, for both proteins, with sufficient sensitivity to apply to model cells and solid tumors. Prior enrichment is required to apply the methods to ORF2p. Although we identified selectivity issues with the *α*-ORF2p antibody used here, the prototype assay presented can now be paired with even more selective enrichment reagents in the pursuit of superior results.

## Methods

### Tissue culturing and cryomilling

The procedures for culturing cells and cryomilling cells to powder has been described in detail previously [81]. In brief, N2102Ep cells were maintained in DMEM, high glucose, GlutaMAX (Gibco, USA) supplemented with 10% (v/v) FBS (Gibco), and 2 mM L-glutamine (Gibco) at 37 °C with 8% CO_2_. Upon reaching 90% confluency they were split 1:5 into progressively larger tissue culture plates and ultimately seeded onto 500 cm2 plates. Once confluent, cells were harvested by physical scraping, washed in PBS, pelleted in a syringe, and extruded directly into a tube containing liquid nitrogen. The frozen cell pellets were cryomilled to a fine cell powder that was used as input material for IP. Resected human tumors and control tissues were cryomilled in the same manner. HEK293TLD cells were used to prepare extracts containing ectopically expressed L1, carried out as previously described [83]. On day 0, four L square glass bottles each containing 200 mL of suspension culture at *∼*2.5 × 10^6^ cell/mL were transfected using 1 µg/mL DNA and 3 µg/mL polyethylene ‘Max’ 40 kDa (Polysciences, Warrington, PA, #24765). The pre-premixing of DNA and PEI-Max was incubated for 20 min at room temperature before added to the bottles. On day 1, cells (200 mL) were split 1:2.5 without changing the medium. On day 3, the cells were induced with 1 µg/ml doxycycline, and on day four the cells were harvested, extruded into liquid nitrogen, and cryomilled as above.

### shRNA-mediated LINE-1 knockdown in N2102Ep

A non-targeting ‘scramble shRNA’ control sequence and two L1-ORF1-targeting shRNAs [9] were cloned into pLKO.1-TRC (Addgene #10878):

- scramble: caacaagatgaagagcaccaactcgagttggtgctcttcatcttgttg
- shRNA1: gaaggcttcagacgatcaactcgagttgatcgtctgaagccttc
- shRNA2: atgaagcgagaagggaagtctcgagacttcccttctcgcttcat

For lentiviral production, one day before transfection, HEK293T cells were seeded at 5.0 × 10^6^ in a 10-cm dish maintained in DMEM, high glucose, GlutaMAX (Gibco, USA) supplemented with 10% (v/v) FBS (Gibco), and 2 mM L-glutamine (Gibco) at 37°C with 5% CO2. Cells were transfected by (1) premixing 30 µL PEI ‘Max’ 40 at 1 mg/mL with 600 µL OptiMEM FBS-free and incubating for 5 min; then (2) this mix was combined with 8.6 µg of shRNA pLKO.1 construct, 2.8 µg of pMD2.G (Addgene #12259) and 8.6 µg of psPAX2 (Addgene #12260) and incubated for 20 min at room temperature; (3) this combined mixture was then added to the cells. Each conditioned medium, containing recombinant lentivirus, was collected after 48 hr and passed through a 0.45 µm filter. The shRNA containing lentivirus-medium was added to N2102Ep cells previously seeded in a 60-mm dish. 48 hr after transduction, the cells were selected with 0.9 µg/mL puromycin (InvivoGen, #ant-pr-1). Puromycin was maintained in all cultures until the control cell cultures were dead (un-treated and treated without pLKO.1). The transduced and selected, shRNA-expressing N2102Ep cells were cultured and cryomilled as described above.

### Immunoprecipitation

Tissue extracts and IPs were produced and conducted as previously described [81]; the extraction solution used in all cases was 20 mM HEPES pH 7.4, 500 mM NaCl, 1% (v/v) Triton X-100, supplemented with protease inhibitors; washing solutions were identical without protease inhibitors. IP samples used for the broad peptide screen in **Figure 2** were prepared using 100 mg HEK293TLD / pLD401 powder and 5 µl (slurry) Dynabeads M-270 Epoxy (ThermoFisher Scientific) conjugated with *α*-ORF2p cl. (MT)9 [24]. For all other MS experiments, IPs were done using 200 mg tissue powder and 20 µl (slurry) Dynabeads M-270 Epoxy conjugated with either *α*-ORF2p cl. 9 or naïve rabbit IgG (mock) control (Innovative Research Inc. #IR-RB-GF-717). Affinity media and clarified extracts were incubated for 1 hr at 4 °C, washed three times with extraction solution and eluted with either 40 mM Tris, 2% (w/v) SDS or 1.1x NuPage LDS loading buffer at 70 °C. For IP-western blotting shown in **Figure 1A**: 10 µl of *α*-ORF1p (4H1, Millipore Sigma #MABC1152) or 10 µl of naïve mouse IgG (mock) control (Sigma #I5381) conjugated Dynabeads M-270 Epoxy media (slurry) were used per 100 mg cell powder. In **Figure 2B**, 20 µl of *α*-ORF2p cl. 9 beads or 20 µl of naïve rabbit IgG (mock) control beads were used per 100 mg cell powder. For IP-western blotting shown in **Figure 4A**: 10 µl (N2102Ep cells) or 5 µl (tumor tissues) *α*-ORF1p magnetic affinity media (slurry) were used per 50 mg powder. All IP-western reactions were incubated for 1 hr at 4 °C, washed three times in extraction solution, and eluted in 1.1x NuPage LDS loading buffer at 70 °C for 5 min.

### Western blotting

LDS eluted IP samples were reduced with dithiothreitol and SDS-PAGE was performed on 4-12% NuPAGE Bis-Tris Mini protein gels (Invitrogen) and wet transferred (0.025% [w/v] SDS / 20% [v/v] methanol in transfer buffer) for 90 min at 70 V, 4 °C onto a PVDF membrane (0.45 µm). Blocking was done for 1 hr at room temperature using 5% (w/v) nonfat dry milk in TBST (20 mM Tris-Cl, 137 mM NaCl, 0.1% [v/v)]Tween 20), pH 7.6. Primary antibodies were applied overnight at 4 °C in 5% (w/v) BSA in TBST, pH 7.6. HRP-conjugated secondary antibodies were applied for 1 hr at room temperature in 5% (w/v) BSA in TBST, pH 7.6. An ImageQuant LAS-4000 system (GE Healthcare) was used for blot imaging on the high sensitivity setting with incremental image capture after the membrane had been treated with chemiluminescent HRP substrate (Millipore Sigma #WBLUF0100). ECL signal capture times displayed varied with target from *∼*1–5min and were free of pixel saturation in any signal displayed in the figures. *α*-ORF1p (4H1, Millipore Sigma #MABC1152) was used at 0.4 µg/ml, *α*-ORF2p cl. 9 was used at 0.7 µg/ml, *α*-ORF2p cl. 5 was used at 2.1 µg/ml, and *α*-ORF2p cl. 11 was used at 1.5 µg/ml. Secondary antibodies *α*-mouse HRP conjugate (Cytiva #NA931) was used at 1:10,000 and *α*-rabbit HRP conjugate (Cytiva #NA934) was used at 1:5,000.

### Targeted MS assay development for ORF1p and ORF2p

A hybrid approach (bioinformatics tools and LC-MS information, described in [84]) was used to select all peptides between 8-25 amino acids and the list was further refined for peptides that are not only well detected in the LC-MS but also provide accurate quantification for the intended protein targets [84–90]. An overview of all considered ORF1p (Uniprot entry Q9UN81) and ORF2p peptides and their properties can be found in **Supp. Table 1**. For ORF2p peptide selection, an additional filter was used: potential locus-specific single amino acid variations were screened against to mitigate the possibility of failing to detect this ultra-rare protein; 75 ORF2p loci (derived from [24]) were aligned and only peptides encoded by more than 65 of them were considered. Crude synthetic peptides (PEPotec peptides with ^13^C^15^N-labeled C-terminal lysines or arginines; Thermo Scientific) were used to optimize the LC-MS settings (charge state, MS2 fragment ions, collision energies, and retention times) for SRM analyses on a triple quadrupole mass spectrometer (TSQ Altis; Thermo Scientific) with a nano-electrospray ion source - more details below (Targeted MS: data collection and analyses); these were mixed with a tryptic digest of HEK293T cell lysate containing ectopically expressed L1 (from pLD401) to assess their suitability to be synthesized as high quality quantitative standards. Ultimately, for ORF1p, the peptides QANVQIQEIQR, LENTLQDIIQENFPNLAR, LSFISEGEIK, and NLEECITR were selected, and for ORF2p, the peptides IFATYSSDK, TAWYWYQNR, QVLSDLQR, and LETIILSK were selected for synthesis (AQUA QuantProHeavy peptides with ^13^C^15^N-labeled C-terminal lysines or arginines, Thermo Scientific). The optimized LC-MS settings for these eight peptides can be found in **Supp. Table 2**. For PRM experiments, data was acquired for the four peptides (two from each ORF1p and ORF2p) that had given the best results in the SRM assay development and experiments were performed on an Orbitrap Exploris 480 (Thermo Scientific) with an EASY-Spray nano-electrospray ion source - more details below (*Targeted MS: data collection and analyses*).

### Targeted MS: sample preparation

Cell extracts, recombinant ORF2p, and *α*-ORF2p IP samples were all used for the targeted quantitative proteomics analyses after in-gel tryptic digestion (Promega #V5111 [SRM] or Roche #3708985001 [PRM]). For SRM experiments, 31.8 µg total protein from HEK293TLD / pLD401 cell extracts were digested, injecting 200 ng of this digest per measurement in L1 peptide linearity and reproducibility experiments (**Supp. Table 3**) and 75 µg protein from a N2102Ep cell extract was used for direct L1 protein quantification (**Fig. 3B**). For PRM experiments, 3 µg of cell extract was used for both the L1 peptide linearity experiment (HEK293TLD / pLD401; **Supp. Table 5**) and for L1 protein quantification in N2012Ep cell extracts (**Fig. 3C**). For the broad PRM L1 peptide screen (**Fig. 2**), 500 ng recombinant ORF2p (described in [49]) was used for each replicate. For all IP samples, entire elutions were processed by in-gel sample preparation. For SRM samples, all proteins were run on NuPAGE 4-12% Bis-Tris gels (Invitrogen Novex) as a single fraction for 5 min at 100 V. The gel was stained with Bio-Safe Coomassie stain (Bio-Rad #1610786) and destained with Milli-Q water. For PRM samples, gels were first run empty for 3 min at 200V and then run with sample for 3 min at 200V. The gel was stained with InstantBlue Coomassie protein stain (Abcam #ab119211) for 1 hr and washed with Milli-Q water overnight. For both SRM and PRM samples, the gel region was sliced into small pieces, washed twice for 30 min at room temperature with mixing at 500 RPM (once with 30% and once with 50% (v/v) aqueous acetonitrile in 100 mM ammonium bicarbonate), and lastly with 100% acetonitrile for 5 min before drying (heat or vacuum). The proteins were reduced with 30 µL of 10 mM dithiothreitol (in 100 mM ammonium bicarbonate) for 30 min at 55 °C and then alkylated with 30 µL of 55 mM iodoacetamide (in 100 mM ammonium bicarbonate) for 30 min, in the dark, at room temperature. The gel pieces were then washed with 100% acetonitrile for 30 min while mixing (500 RPM) and dried before overnight digestion with 30 µL trypsin. For SRM, 10 ng/µL of trypsin was used for IP samples and 16.7 ng/µL of trypsin was used for the cell lysates; for PRM 12.5 ng/µl of trypsin was used for all samples. Digestion was carried out at 37 °C, overnight; the next day, the peptides were eluted from the gel pieces with 75% (v/v) acetonitrile, 5% (v/v) formic acid. The eluted fractions were dried under vacuum and resuspended in 0.1% (v/v) formic acid spiked with heavy labeled AQUA QuantPro peptides (Thermo Scientific) to a final volume of 20 µL. For targeted experiments, 2-4 µL of sample was injected per measurement.

### Targeted MS: data collection and analyses

#### SRM

Chromatographic separation of the peptides was performed by liquid chromatography on a nano-UHPLC system (Ultimate UHPLC focused; Dionex) using a nano column (Acclaim PepMap 100 C18, 75 µm x 50 cm, 2 µm, 100 Å; Dionex). Samples were injected using the µL-pickup system with 0.1% (v/v) formic acid as a transport liquid from a cooled autosampler (5 ^*°*^C) and loaded onto a trap column (µPrecolumn cartridge, Acclaim PepMap100 C18, 5 µm, 100 Å, 300 µm id, 5 mm Dionex). Peptides were separated on the nano-LC column using a linear gradient from 3-40% (v/v) acetonitrile plus 0.1% (v/v) formic acid in 90 min at a flow rate of 300 nL/min. The mass spectrometer was operated in the positive mode at a spray voltage of 2000 V, a capillary temperature of 275 ^*°*^C, a half maximum peak width of 0.7 for Q1 and Q3 and a cycle time of 1 msec. The measurements were scheduled in windows of 10 minutes around the predetermined retention time (targeted m/z list can be found in **Supp. Table 2**). *PRM* : Chromatographic separation of the peptides was performed by liquid chromatography on an EASY-nLC 1200 system (Thermo Scientific) using a nano-column (PepMap RSLC C18, 75 µm x 50cm, 2 µm, 100 Å) with a temperature maintained at 50 °C. Samples were injected using the µL-pickup system using buffer A (0.1% (v/v) formic acid) as a transport liquid from a cooled autosampler (7°C) and loaded directly onto the analytical column. Peptides were separated using a linear gradient from 3-29% buffer B (95% (v/v) acetonitrile, 0.1% (v/v) formic acid) in 87 minutes at a flow rate of 250 nL/min. The mass spectrometer was operated in PRM mode selecting precursor m/z for fragmentation corresponding to ORF1p/ORF2p peptides and heavy labeled equivalents (targeted m/z list can be found in **Supp. Table 2**). The instrument was operated with a spray voltage of 1800 V, a capillary temperature of 305 °C. A precursor MS scan (m/z 300-1000, positive polarity) was acquired at a nominal resolution of 120K, followed by HCD-MS/MS at a nominal resolution of 15K. HCD was performed using stepped collision energies of 25/30/35% and normalized AGC target was set to 100% with a maximum injection time of 100ms. For both SRM and PRM data, LC-MS peak assignments were manually curated using the Skyline software. The sum of peak areas from all transitions (see Supp. **Table 2**) for the endogenous and isotopically-labeled standard peptides was used to calculate the ratio between the endogenous and standard peptides. The known amount of the isotopically-labeled standard peptides were used to calculate the amount of the endogenous peptide. Endogenous ORF2p abundance calculation: 200 mg N2102Ep cells were used per experiment, corresponding to *∼*20 mil cells. Following work-up, 10% of the resulting sample was injected for analysis by targeted LC-MS/MS, resulting in an average detection of *∼*70 amol ORF2p (**Fig. 3D**) – providing for the following estimation: (7 × 10^−17^ mol ORF2p/measurement * 6.023 × 10^23^ molecules/mol) / 20 mil cells = 2.1 ORF2p molecules per cell, or 21 molecules corrected for the fraction of sample analyzed by MS (10%). Further correcting for losses in sample preparation, of up to *∼*60% [84], yields a likely average range of 20–50 ORF2p molecules per cell.

### Shotgun MS and Data Processing of *α*-ORF2p IPs

For shotgun proteomic analyses, IP elutions were processed using S-Trap columns [91]. The dried-down samples were resuspended in 25 µL at a final concentration of 5% (w/v) SDS, 8 M urea, 100 mM glycine, pH 7.55 with MMTS used to block cysteine residues. Samples were incubated with a Trypsin/Lys-C mix (Promega #V5071) for 1 hr at 47 °C. the peptides eluted from S-Trap columns were dried and resuspended in 25 µl water:methanol:formic acid solution (94.9:5.0:0.1 parts by volume). For experiments using DDA mode, 5 µl of the final sample was loaded onto a 75 µm x 50 cm Acclaim PepMap™ RSLC nano Viper column filled with 2 µm C18 particles (Thermo Fisher Scientific, Bremen, Germany) via a Dionex Ultimate™ 3000 HPLC system interfaced with an Orbitrap Exploris™ 480 mass spectrometer (Thermo Fisher Scientific). Column temperature was set to 35 °C. Using a flow rate of 300 nl/min, peptides were eluted in a gradient of increasing acetonitrile, where solvent A is 0.1% formic acid in water and solvent B is 0.1% formic acid in acetonitrile. Peptides were ionized by electrospray at 2 kV and eluted over a 60 min gradient (3% B over 3 min; 3 to 50% B over 45 min; 2 min to 80% B; then wash at 80% B over 5 min, 80 to 3% over 2 min and then the column was equilibrated with 3% B for 3 min). Full scans were acquired in profile mode (m/z 200-2000, positive polarity) at 120,000 nominal resolution and the top 25 most intense precursor ions with charge states +2-6 in each scan were fragmented by HCD at 30% normalized collision energy. Precursors previously sequenced were excluded for 20 s, within a mass tolerance of 10 ppm. Fragmentation spectra were acquired in centroid mode with resolution at 15,000. The normalized AGC target was set to 300% with a maximum injection time of 50 ms. For experiments using DIA mode (high resolution MS1 (HRMS1) method [56, 92]), 5 µl sample was loaded onto a 75 µm x 40 cm Acclaim PepMap™ RSLC nano Viper column (1.9 µm C18 particles; Thermo Fisher Scientific), plumbed to a Dionex Ultimate™ 3000 RSLC system interfaced with an Orbitrap Exploris™ 480 mass spectrometer (Thermo Fisher Scientific). The column temperature was set to 40 °C and the flow rate was set to 300 nl/min. The peptides were eluted in a gradient of increasing acetonitrile, where solvent A was 0.1% (v/v) formic acid in water and solvent B was 0.1% (v/v) formic acid in acetonitrile. The peptides were separated using a 120 min gradient (2% B over 3 minutes; 3% to 45% B over 87 min; 1 min to 80%; then wash at 80% over 13 min; 80% to 2% over 1 min and then the column was equilibrated with 2% B for 16 min); eluting peptides were ionized by electrospray at 2 kV. The full MS scans were acquired (m/z 400-1200, positive polarity) at a nominal resolution of 120,000. The precursor ions were isolated within a 8.6 Da window with an m/z range of 400 and 1198; these were fragmented by HCD using normalized collision energy of 32%. The normalized AGC target was set to 1000%.

For the DDA shotgun proteomics, a full analysis pipeline has been described by [93]. In brief, the raw files were processed in MaxQuant (version 2.3.1) [94] and searched against a human-specific proteomic database (UniProt, February 2024, containing 20360 canonical entries) supplemented with ORF1p and ORF2p non-redundant sequences [24]. The default settings were used with the exception of methylthio modification of cysteines as a fixed modification (from MMTS), and both match between runs and IBAQ were enabled. From the protein groups file, those marked as ‘contaminants’ or ‘reverse’ by MaxQuant were removed. Only proteins that had ‘Peptide counts (razor+unique)’ *≥*2 were retained for analysis. Intensities of the proteins MAGOH and MAGOHB were summed together in every experiment. Protein LFQ intensities were log2-transformed and LFQ missing intensity values were imputed as previously described [93]. For statistical testing, the log2-transformed intensities were used and proteins were subjected to unpaired, two-sample t-test between ORF2p IPs and rabbit IgG (mock control) IPs (four replicates in each condition). Proteins co-enriched with ORF2p were considered statistically significant if the Benjamini-Hochberg adjusted p-values were ¡ 0.05 and log2 fold-change was *≥*1.

For DIA shotgun proteomics, the RAW data collected from DIA experiments were processed using Spectronaut [95] and searched against a human-specific proteomic database (UniProt, February 2024, containing 20360 canonical entries). Methylthio modifications to cysteines (from MMTS) were specified in the Pulsar Search Settings and all other modification options were left as default. The protein groups file was exported and the same data processing approach was used as in the DDA experiments. Statistical significance and fold-changes were calculated in R (version 4.1.2) using the bioconductor package “Limma” [96] (version 3.50.3) using the log-transformed values: the scramble shRNA control samples were compared to the two distinct L1 shRNA KD samples (combined as one group). Proteins were considered to be significantly changed with adjusted p-value *≤*0.05 and log_2_ fold-change *≥* 1. iBAQ values from the significant proteins were used to calculate the relative iBAQ (riBAQ [97]), which is the iBAQ for a given protein divided by the total iBAQ generated by the sample; to present the data in terms protein copy number compared with ORF1p, ORF1p iBAQ was set to 1.

#### Supplementary information

This manuscript is supplemented by eleven tables which are collated in the accompanying file: *Supp Tables*.*xlsx*

#### Availability of data and materials

All the MS data files have been deposited to the ProteomeXchange Consortium [98] via the PRIDE [99] partner repository with the data set identifier PXD057269. SRM data has also been submitted to the PASSEL database under the identifier PASS05852.

#### Ethics approval

Fresh frozen ovarian tumors were collected at Massachusetts General Hospital Department of Pathology as de-identified patient samples in accordance with Exemption 4, of research involving human subjects, from the National Institutes of Health. These samples were subsequently analyzed at The Rockefeller University, where, according to 45 CFR 46.102 (f) of the U.S. Dept. of Health and Human Services, it was determined that this research does not involve human subjects (IRB reference #334332).

#### Competing interests

JL reports grants, personal fees, and equity from Rome Therapeutics, outside the submitted work. MT reports personal fees, and equity from Rome Therapeutics, outside the submitted work. JL, MIN., and JCW. have a patent application pending, based on this work. The other authors declare no competing interests.

#### Authors’ contributions

Conceptualization: JL; Data Curation: MIN, JCW, OGRB, AM; Formal analysis: MIN, JCW, OGRB; Funding acquisition: JL; Investigation: MIN, JCW, HJ, LHDS, MO, SH, MJN, AvP, MS, LtM, AM; Methodology: MIN, JCW, JL; Project administration: JL; Resources: JCW, VD, MT, JL; Software: OGRB, MO; Supervision: JCW, JL; Validation: MIN, JCW, MJN; Visualization: MIN, OGRB, AM, JL; Writing - Original Draft: MIN, JCW, JL.

## Acknowledgments

We thank Profs. Brian T. Chait and Michael P. Rout for providing NCDIR resources and research infrastructure, used in-part to facilitate the PRM aspects of this study. We thank Kelly R. Molloy for carrying out GeLC-MS on an ORF2p IP. We thank the IMSC in Groningen for the infrastructure to facilitate the development and application of the SRM experiments.

## Funding

This work was funded in-part by the National Institutes of Health (NIH) grants R01GM126170, R01AG078925, and R01AI186337, The Robertson Therapeutic Development Fund (through The Rockefeller University), and the UMCG Kanker Researchfonds, to JL. This project also benefited from support by the National Center for Dynamic Interactome Research, funded by NIH grant P41GM109824, and received administrative support from the National Center for Advancing Translational Sciences (through The Rockefeller University), funded by NIH grant UL1TR001866.

## Notes

### Competing Interest Statement

JL reports grants, personal fees, and equity from Rome Therapeutics, outside the submitted 550 work. MT reports personal fees, and equity from Rome Therapeutics, outside the submitted work. JL, MIN., and 551 JCW. have a patent application pending, based on this work. The other authors declare no competing interests.

